# Do chemical language models provide a better compound representation?

**DOI:** 10.1101/2023.11.07.566025

**Authors:** Mirko Torrisi, Saeid Asadollahi, Antonio de la Vega de León, Kai Wang, Wilbert Copeland

## Abstract

In recent years, several chemical language models have been developed, inspired by the success of protein language models and advancements in natural language processing. In this study, we explore whether pre-training a chemical language model on billion-scale compound datasets, such as Enamine and ZINC20, can lead to improved compound representation in the drug space. We compare the learned representations of these models with the de facto standard compound representation, and evaluate their potential application in drug discovery and development by benchmarking them on biophysics, physiology, and physical chemistry datasets. Our findings suggest that the conventional masked language modeling approach on these extensive pre-training datasets is insufficient in enhancing compound representations. This highlights the need for additional physicochemical inductive bias in the modeling beyond scaling the dataset size.

## 1 Introduction

Language models have revolutionized the field of natural language processing, and protein representation [11, 17, 22]. This has stimulated the development of numerous chemical language models (CLMs) [5, 8, 10, 26], on increasingly large portions of public compound databases, such as ChEMBL [16] and ZINC20 [9]. CLMs are pre-trained using masked language modeling, and then fine-tuned on down-stream tasks of interest, often resulting in state-of-the-art predictive performance [2, 23].

The availability of multi-billion scale compound databases, such as ZINC20 [9] and Enamine [21], prompted us to investigate whether pre-training a CLM on billion-scale compound datasets can suffice to outperform the de facto standard compound representation for drug discovery, i.e., Extended-Connectivity Fingerprints (ECFP) [19]. Additionally, this study is the first to investigate the potential impact of using the Enamine REAL Database for pre-training CLMs, as most recent studies have focused on exploiting ZINC20 or ChEMBL.

Our investigation aims to explore the effectiveness of pre-training CLMs on various compound databases of up to 2 billion compounds. We evaluate whether scaling the pre-training dataset can improve the ability of CLMs to represent the drug space compared to ECFP. To achieve this, we benchmark the predictive performance of these representations on biophysics, physiology, and physical chemistry datasets. By conducting these experiments, we aim to determine whether CLMs can provide a richer representation of the drug space than ECFP without fine-tuning.

## 2 Methods

### 2.1 Datasets standardization

All compounds in both pre-training and benchmarking datasets are originally represented using SMILES. We standardize all SMILES using DataMol 0.10 [15], and encode them as SELFIES using selfies 2.1.1 [12]. We opt for using SELFIES to facilitate the learning process, since it was specifically developed for generative modeling and provides a guarantee of molecular validity. We remove all compounds that do not pass the standardization procedure, or are not encodable as SELFIES, or are found to be duplicates.

#### 2.1.1 Pre-training datasets

The pre-training datasets in this study are derived from 3 public compound databases: ChEMBL 33 [16], Enamine REAL Database [21], and ZINC20 [9]. After being standardized, ChEMBL, Enamine, and ZINC20 are described by 341, 67, and 277 distinct SELFIES symbols, respectively (see Table 1).

**Table 1:** SELFIES symbols shared by pre-training and benchmark datasets.

| Name | Pre-training datasets |  |  |  | Tot |
| --- | --- | --- | --- | --- | --- |
|  | ChEMBL | Enamine | ZINC20 | All |  |
| MoleculeNet | 147 | 60 | 108 | 56 | 224 |
| Papyrus1k | 71 | 35 | 60 | 35 | 73 |
| Tot | 341 | 67 | 277 |  |  |

The standardized databases are split assigning 99%, 0.5%, and 0.5% of the compounds to the training, validation, and test set, respectively. We obtain ChEMBL-2M splitting ChEMBL at random. Enamine- 2M and ZINC20-2M are randomly sampled from Enamine and ZINC20, respectively, while keeping the same amount of compounds in their training, validation, and test sets as in ChEMBL-2M.

Similarly, we obtain ZINC20-2B splitting ZINC20 at random, and Enamine-2B sampling from Enamine at random (i.e., about 1.9*B* compounds in total). The size of each dataset split is reported in Table 2.

**Table 2:**
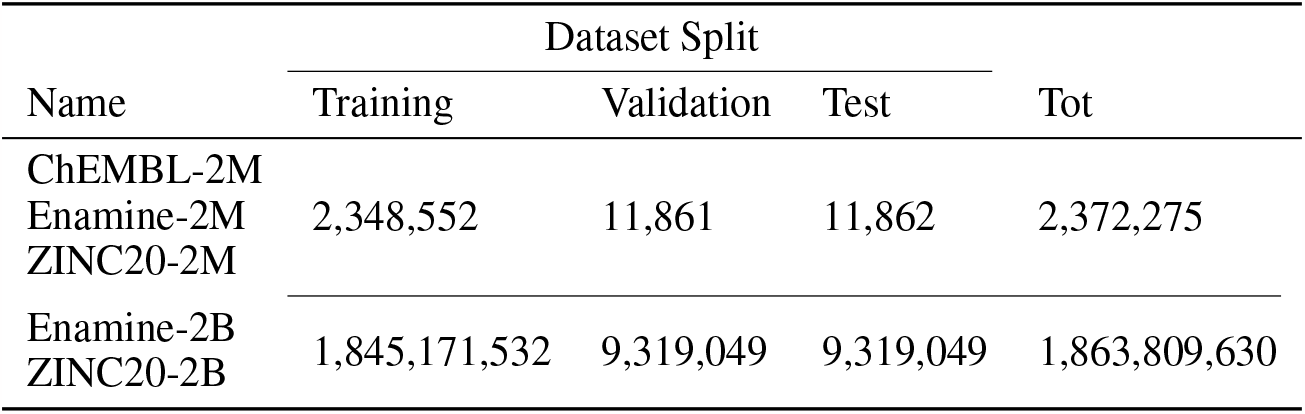
Pre-training datasets: number of compounds per dataset split.

#### 2.1.2 Papyrus1k

Papyrus1k is derived from Papyrus, a public database consisting of nearly 60*M* compound-protein pairs, 1.2*M* of which are considered of high quality, i.e., “representing exact bioactivity values measured and associated with a single protein or complex subunit” [4]. We focus on these high quality pairs, and keep only those involving proteins observable in at least 1*k* bioactivities. We binarize all bioactivities by setting a *pIC*_50_ threshold value of 6.

After standardizing and encoding all compounds as SELFIES, we find 73 distinct SELFIES symbols in Papyrus1k. Out of these, 35 symbols are found in all pre-training datasets (see Table 1). Thus, we remove 1,698 compounds (4,031 bioactivities) containing SELFIES symbols not found in at least one pre-training dataset. We randomly split compounds per protein, assigning 80%, 10%, and 10% for training, testing, and validation, respectively. The final benchmark dataset contains 489,402 compounds, 280 proteins, and 881,081 bioactivities (see Table 3).

**Table 3:**
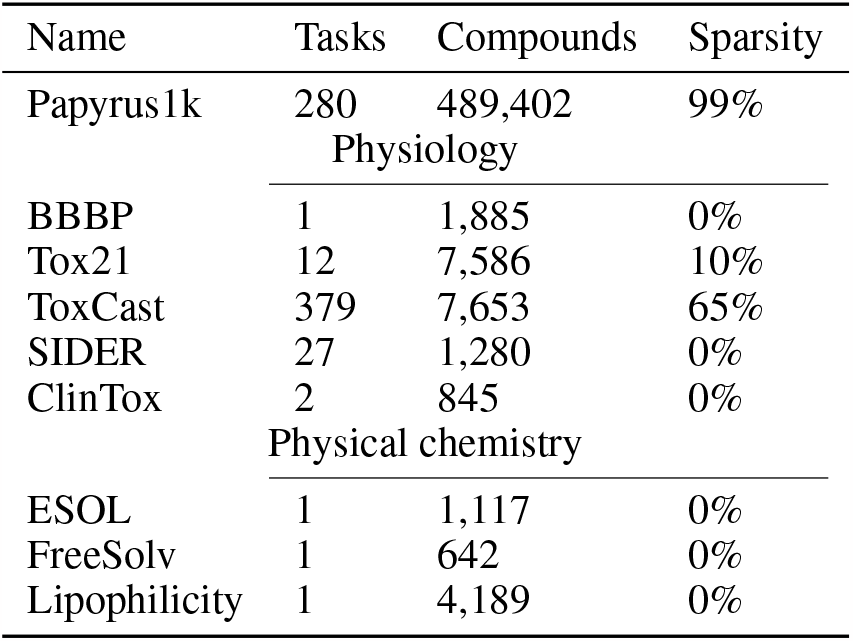
Number of unique compounds and tasks in the benchmark datasets, and their sparsity.

| Name | Tasks | Compounds | Sparsity |
| --- | --- | --- | --- |
| Papyrus1k | 280 | 489,402 | 99% |
| Physiology |  |  |  |
| BBBP | 1 | 1,885 | 0% |
| Tox21 | 12 | 7,586 | 10% |
| ToxCast | 379 | 7,653 | 65% |
| SIDER | 27 | 1,280 | 0% |
| ClinTox | 2 | 845 | 0% |
| Physical chemistry |  |  |  |
| ESOL | 1 | 1,117 | 0% |
| FreeSolv | 1 | 642 | 0% |
| Lipophilicity | 1 | 4,189 | 0% |

#### 2.1.3 MoleculeNet

MoleculeNet [25] is a collection of benchmark datasets for assessing machine learning methods for characterizing compounds. To expand into more areas related to drug discovery and development, we focus on two categories of the datasets in MoleculeNet: physiology and physical chemistry. We do not include the biophysics datasets because of their similarity to Papyrus1k, and the quantum mechanism datasets as we feel it is less related to drug discovery. The physical chemistry datasets are the only regression datasets in this study.

After standardizing, removing duplicates, and encoding the compounds in these datasets as SELFIES, we find 224 distinct SELFIES symbols. Out of these, only 56 are in the pre-training datasets (see Table 1), yet describe 17,329 compounds (25,197 pairs). We, thus, remove 1,787 compounds (3,275 pairs) for containing SELFIES symbols not found in at least one pre-training dataset. We also remove 3 compounds for requiring more than 512 symbols, i.e., the maximum supported length by our CLMs. The composition of the final benchmark datasets, and their sparsity, is in Table 3.

For ToxCast, i.e., one of the physiology datasets, we keep only 379 tasks (out 617) associated to at least 1*k* compounds. For each dataset, we randomly split compounds per task, assigning 80%, 10%, and 10% for training, testing, and validation, respectively.

### 2.2 Chemical language modeling

To construct the character-based tokenizer and the CLM architecture, specifically Bert-Large [6], we employ Transformers 4.28.1 [24]. We opt for Bert-Large as our model architecture because it is recognized as a ubiquitous baseline for masked language modeling [18]. We utilize Lightning 2.0.1 [7] to facilitate multi-node training on 2 NVIDIA DGX-A100 machines.

We pre-train all CLMs with masked language modeling (15%), Adam optimizer (*β*_1_ = 0.9, *β*_2_ = 0.999, and *E* = 1*e−* 8), learning rate of 5*e −* 5, weight decay set to 0.1, and a cosine scheduler. We set the warm up steps to 1*k* and the epoch to 1 for ZINC20-2B and Enamine-2B. We reduce the warm up steps to 100 and increase the epochs to 5 for ChEMBL-2M, Enamine-2M, and ZINC20-2M.

### 2.3 Benchmark modeling

We utilize Scikit-learn 1.2.2 [3] to train either a Random Forest (RF) classifier or regressor for each task in the benchmark datasets. Each RF is trained on either ECFP or the average pool of the last hidden layer of a CLM, i.e., the CLM embedding.

To generate traditional fingerprints, we employ the Morgan fingerprints algorithm in RDKit 2022.9.5 [13] with 2, 048 bits and radius 0, 1, or 2, corresponding to ECFP0, ECFP2, and ECFP4, respectively. For generating CLM embeddings, we use the final model checkpoint for each CLM.

To evaluate 30 combinations of hyperparameters, we conduct a Bayesian hyperparameter optimization using Optuna [1]. We guide the optimization process using either the Matthew correlation coefficient (MCC), or the root-mean-square error (RMSE) as the evaluation metric. Specifically, we let Optuna suggests the number of estimators (ranging from 100 to 5k), the depth (ranging from 5 to 100), and the number of features to consider (“sqrt” and “log2”).

## 3 Results

We successfully pre-train a CLM model for each pre-training dataset, except for ZINC20-2M, which did not converge. It is worth noting that while all validation losses approach zero, the losses of Enamine-2B and ZINC20-2B, i.e., the CLMs pre-trained on 2*B* compounds, are lower yet continue to decrease (see Figure 1). Previous studies have used similar validation loss values as an indication of effective pre-training [20, 14].

**Figure 1.**
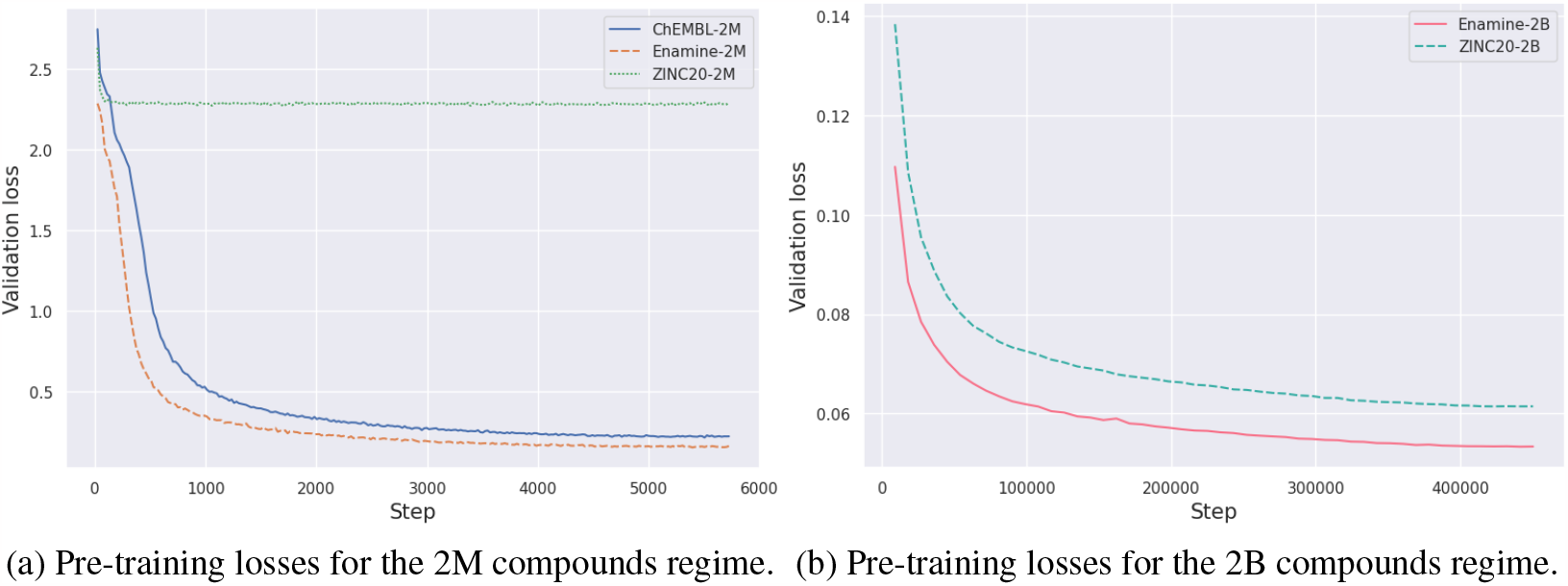
Pre-training validation losses for (a) 2M compounds regime, and (b) 2B compounds regime. The pre-training for (a) lasts for 5 epochs, while for (b) it only lasts for one epoch.

In the following 2 sections, we present benchmark results for CLM embeddings obtained through pre-training on ChEMBL, Enamine, and ZINC20. Surprisingly, we find that CLM embeddings perform similarly, with no clearly noticeable difference scaling the pre-training dataset from 2*M* to 2*B* compounds. We also find that CLM embeddings perform worse than ECFP4 on Papyrus1k, our benchmark dataset of high quality bioactivities (see Section 3.1). Finally, we observe more on par performance on MoleculeNet (see Section 3.2).

To further investigate why CLMs do not provide a better compound representation than ECFP4, we report results for ECFPs with smaller radius. A ECFP with radius 0 represents the initial atom identifier and simple physicochemical properties of that atom (ECFP0), with radius 1 includes the information of the immediate neighbors (ECFP2), and with radius 2 encompasses the neighbors up to 2 bonds away (ECFP4). Notably, we find that ECFP0 obtains comparable predictive performance to CLMs on both Papyrus1k and MoleculeNet. We also observe that larger radius ECFPs achieves superior predictive performance, especially on Papyrus1k. This suggests that CLMs may not capture the topological information that is beneficial for biophysical applications, which is represented by larger radius ECFPs. Analogous observations can be made on MoleculeNet, although a larger radius seems to be not always advantageous.

### 3.1 Biophysics benchmark

The benchmark results on Papyrus1k show similar results for all compound representations, with ECFP2 and ECFP4 showing an advantage over both ECFP0 and CLM embeddings (see Figure 2). Interestingly, ECFP0 seems to perform on par with CLM embeddings, with a Pearson correlation coefficient of at least 0.86 (see Figure 3a). This may suggest that CLMs are not capturing topological information, and that neighbors up to 1 bond away may provide sufficient biophysics information.

**Figure 2.**
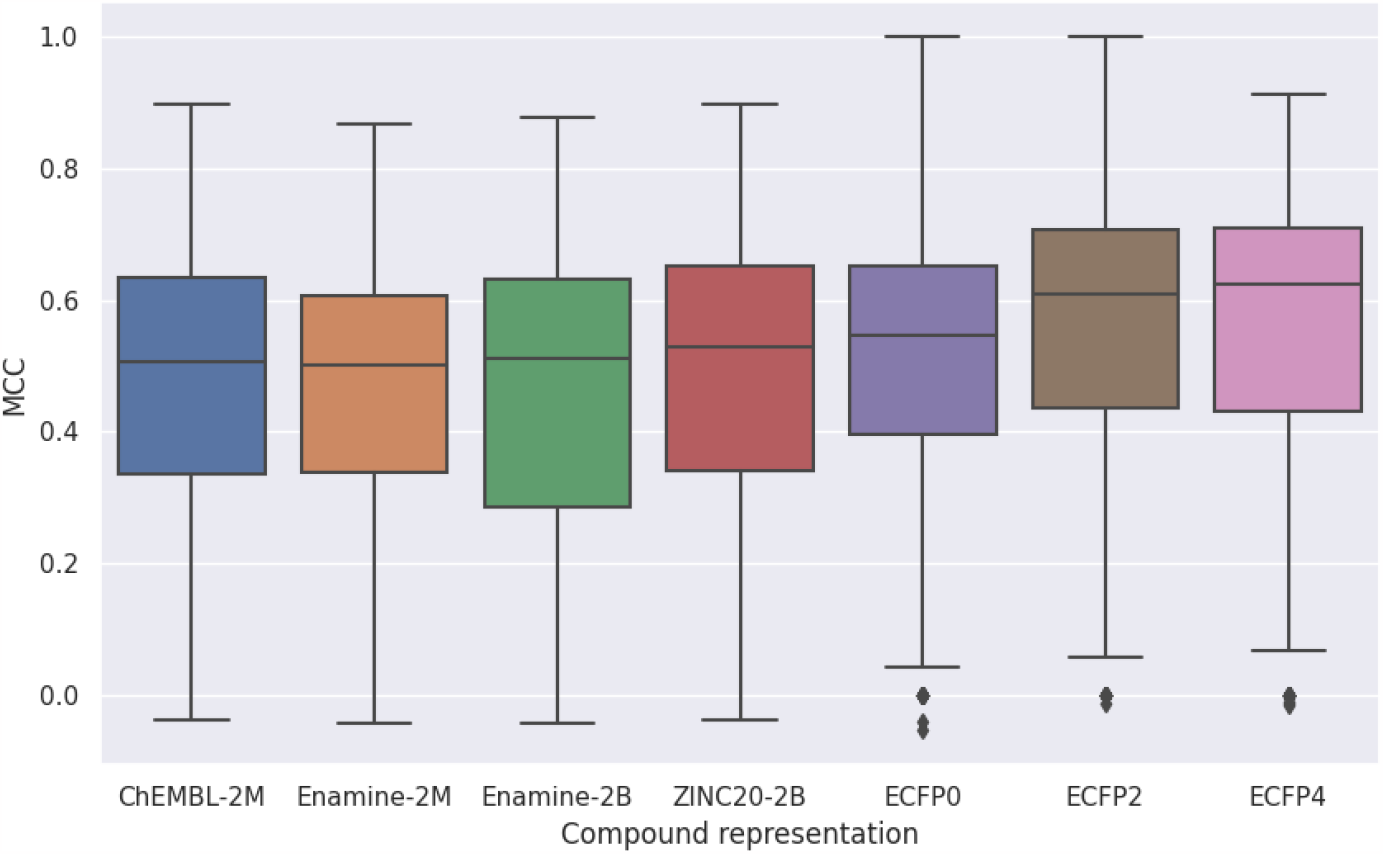
MCC on the high quality test set of Papyrus1k. Each box plot represents the distribution of individual MCC values for the test set of all proteins in the Papyrus1k dataset. ECFP0 seems comparable to CLM embeddings, while ECFP2 seems to provide sufficient biophysics information.

**Figure 3.**
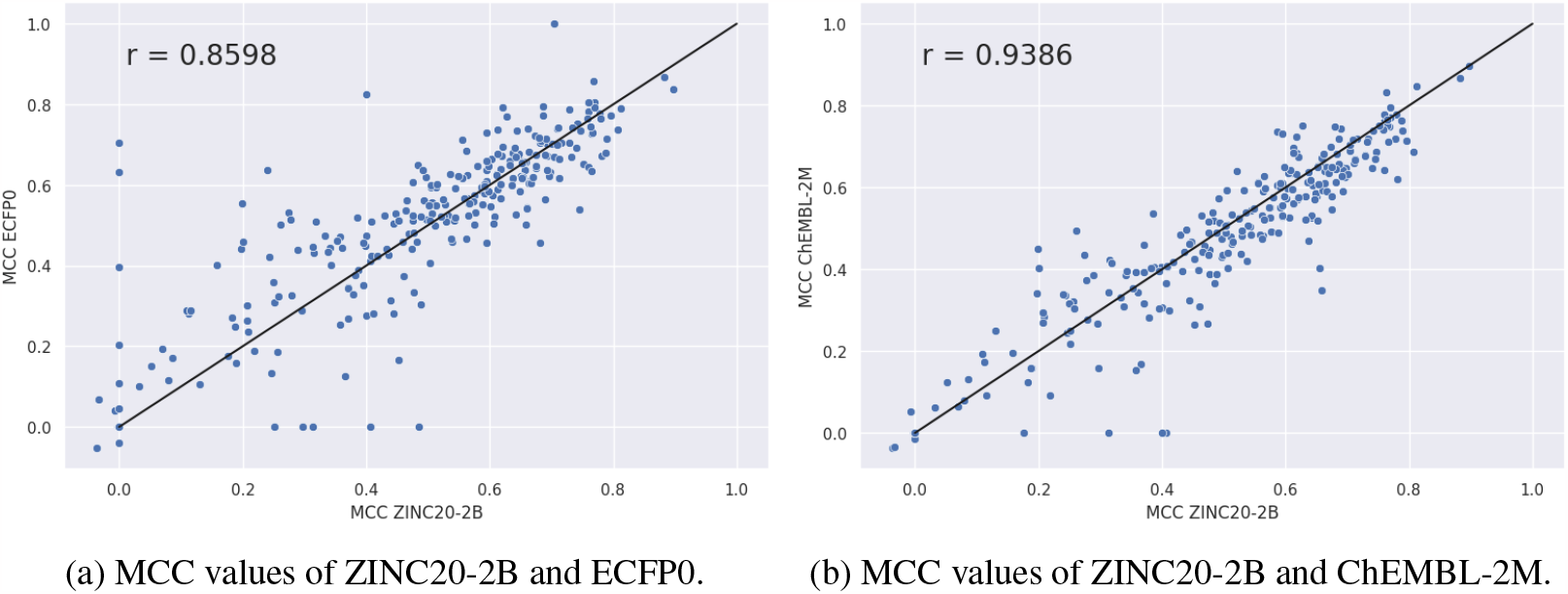
Comparison of two compound representations at a time, and their Pearson correlation coefficient: (a) shows the minimum agreement between a CLM embedding and ECFP0, while (b) an even stronger agreement between CLMs pre-trained on 2*M* or 2*B* compounds.

Moreover, CLMs show an even greater correlation among themselves (see Figure 3b), with no clearly noticeable difference between CLMs pre-trained on 2*M* or 2*B* compounds. This finding may suggest that masked language modeling of Bert-Large on 2*M* compounds suffice to capture atomic identifiers, but may not capture additional valuable insights.

Finally, removing from the pre-training set of ChEMBL-2M the compounds in the validation and test set of Papyrus1k does not appear to affect the benchmark results (results not shown).

The RF hyperparameters and validation MCC are reported in the Supplementary material (see Figure 5).

### 3.2 Physiology and physical chemistry benchmark

The predictive performance of ECFPs on MoleculeNet is somewhat comparable to CLM embeddings, with a less clear benefit from neighbor information compared to Papyrus1k (see Figure 4). Moreover, Enamine-2B appears to be on par with ZINC20-2B in terms of predictive performance for physiology (classification) datasets, while exhibiting a slight disadvantage for physical chemistry (regression) datasets. Overall, ChEMBL-2M seems to provide the best CLM embeddings, which may be attributed to the larger set of SELFIES symbols it contains.

**Figure 4.**
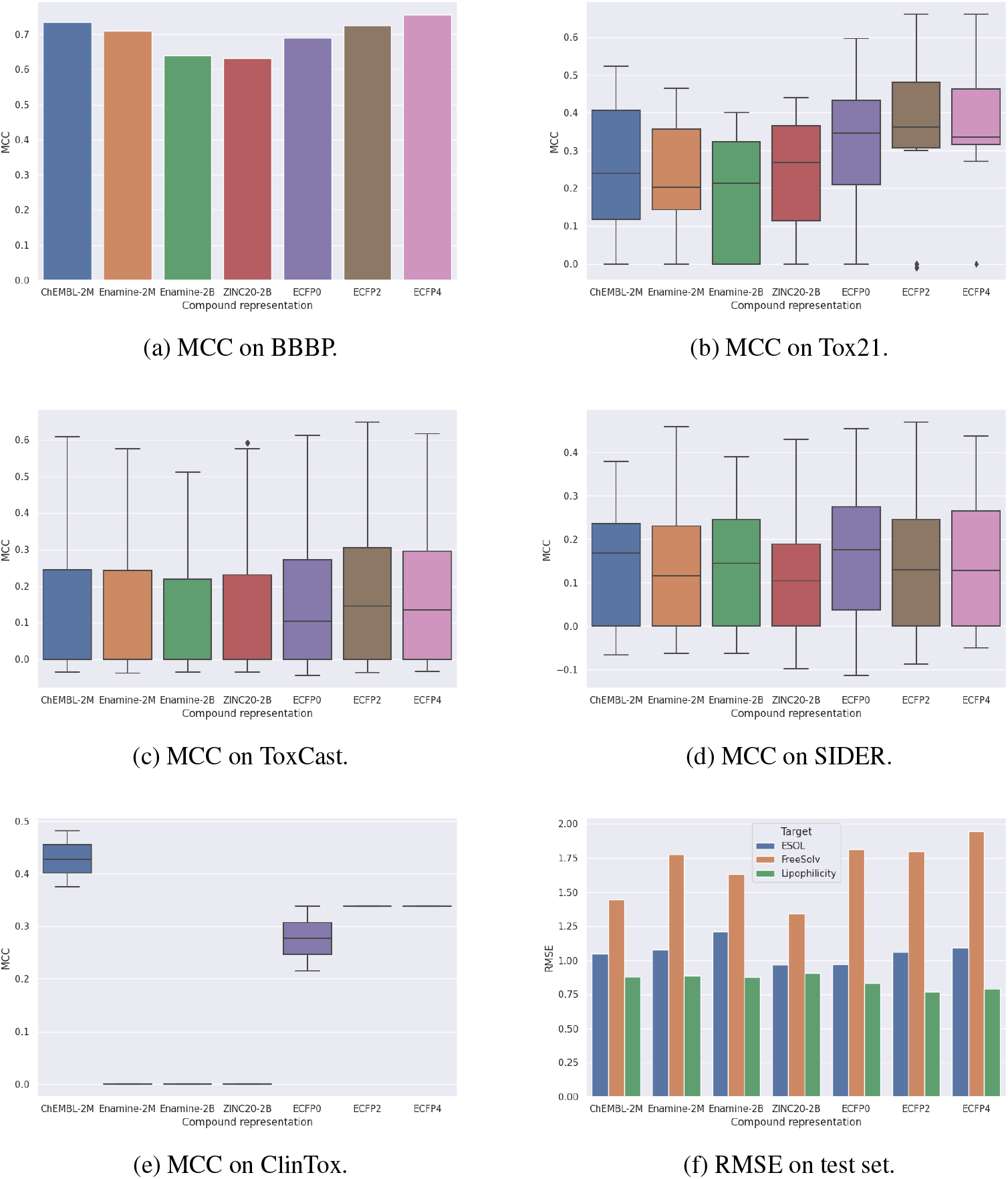
MCC on (a-e) each physiology dataset, and (f) RMSE physical chemistry datasets of MoleculeNet. No clear superiority for ECFPs or CLMs is observable on (a), (d), (e) and (f).

BBBP (binary labels of blood-brain barrier penetration) is the only physiology dataset with a single task, similarly to all physical chemistry (regression) datasets. All compound representations show comparably strong predictive performance, with ChEMBL-2M and Enamine-2M having a slight advantage over Enamine-2B and ZINC20-2B. BBBP is the only MoleculeNet dataset exhibiting some correlation between radius size and predictive performance for ECFPs.

Tox21 (qualitative toxicity measurements on 12 proteins) and ToxCast (toxicology data on 379 tasks) lead to similarly low predictive performance for all compound representations. ECFPs show a slight advantage with increasing radius when looking at the median MCC. Similarly to all physiology datasets except BBBP, ECFP2 performs at least as well as ECFP4.

SIDER (marketed drugs and adverse drug reactions on 27 system organ classes) exhibits low predictive performance for all compound representations. Though, ECFP0 performs slightly better than any other compound representation.

ClinTox (qualitative data of drugs approved and of those that failed clinical trials for toxicity) is the only physiology dataset with less than 1*k* compounds per task. All CLMs leads to 0 MCC except for ChEMBL-2M, which outperforms all compound representations in terms of predictive performance.

This result may indicate that Enamine-2B and ZINC20-2B may have limited generalizability to the broader drug space represented in ClinTox (and ChEMBL).

ESOL (water solubility data for 1*k* compounds) leads to generally strong regression performance, with ECFP0 performing on par with ZINC20-2B and slightly ahead of the remaining compound representation.

FreeSolv (hydration free energy for 642 compounds) is the only physical chemistry dataset where CLM embeddings exhibit superior predictive performance compared to ECFPs. Nonetheless, increasing the maximum depth and number of estimators for the RFs may further enhance the predictive performance on this dataset (see Figure 7 in the Supplementary material).

Lipophilicity (octanol/water distribution coefficient for 4*k* compounds) is the only physical chemistry dataset where ECFP2 and ECFP4 exhibit slightly better performance compared to all other compound representation.

## 4 Conclusion

In this study, we explore whether CLMs yield representations that facilitate accurate predictions on the physical, physiological, and biophysical characteristics of small drug-like molecules. We systematically assess the impact of pre-training data origin and size. Our results show that simply scaling the pre-training of CLMs from millions to billions of example compounds does not lead to molecular representations that outperform ECFP4 on downstream drug prediction tasks. Furthermore, we do not observe substantial differences in predictive performance of CLMs when pre-trained on distinct biomolecule databases, including ChEMBL 33, Enamine REAL Database, and ZINC20. The only notable exception to these observations was the CLM trained on ChEMBL, which performed better than both ECFP4 and other CLMs on a single physiology benchmark dataset.

We further observed that, for most tests, CLMs – regardless of size or origin of the pre-training dataset – appear to have predictive performance comparable to ECFP with a radius of zero. ECFP0 encodes only basic physicochemical properties and the initial atom identifier, while ECFP4 includes topological information in addition to these properties. Based on these observations, we hypothesize that CLMs may not comprehensively learn topological information through masked language modeling. To address this limitation, we propose that future experiments with CLMs applied towards drug discovery and development should explore strategies that embed additional physicochemical inductive biases into the learning procedure.

## Acknowledgments and Disclosure of Funding

The authors are grateful to Remco Loos, Giorgio Tamo, and Matthew Trotter from BMS for useful comments and suggestions.

Antonio de la Vega de Leon is funded thanks to project PTQ2021-011654 by MCIN/AEI/10.13039/501100011033.

## 5 Supplementary material

**Figure 5.**
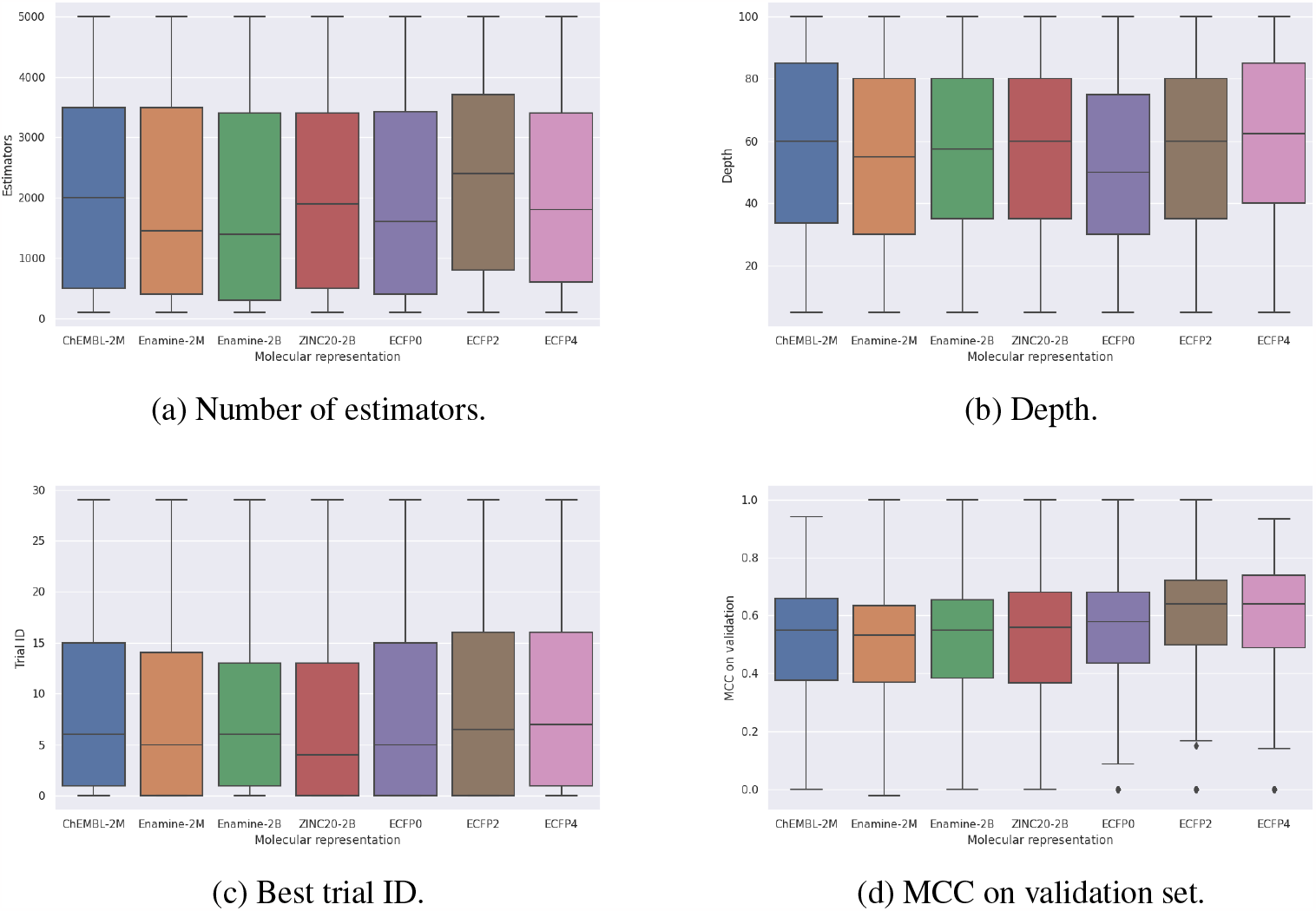
Overview of the best hyperparameters for Papyrus1k (a-b), the number of trials to find them (c), and their MCC on validation set (d).

**Figure 6.**
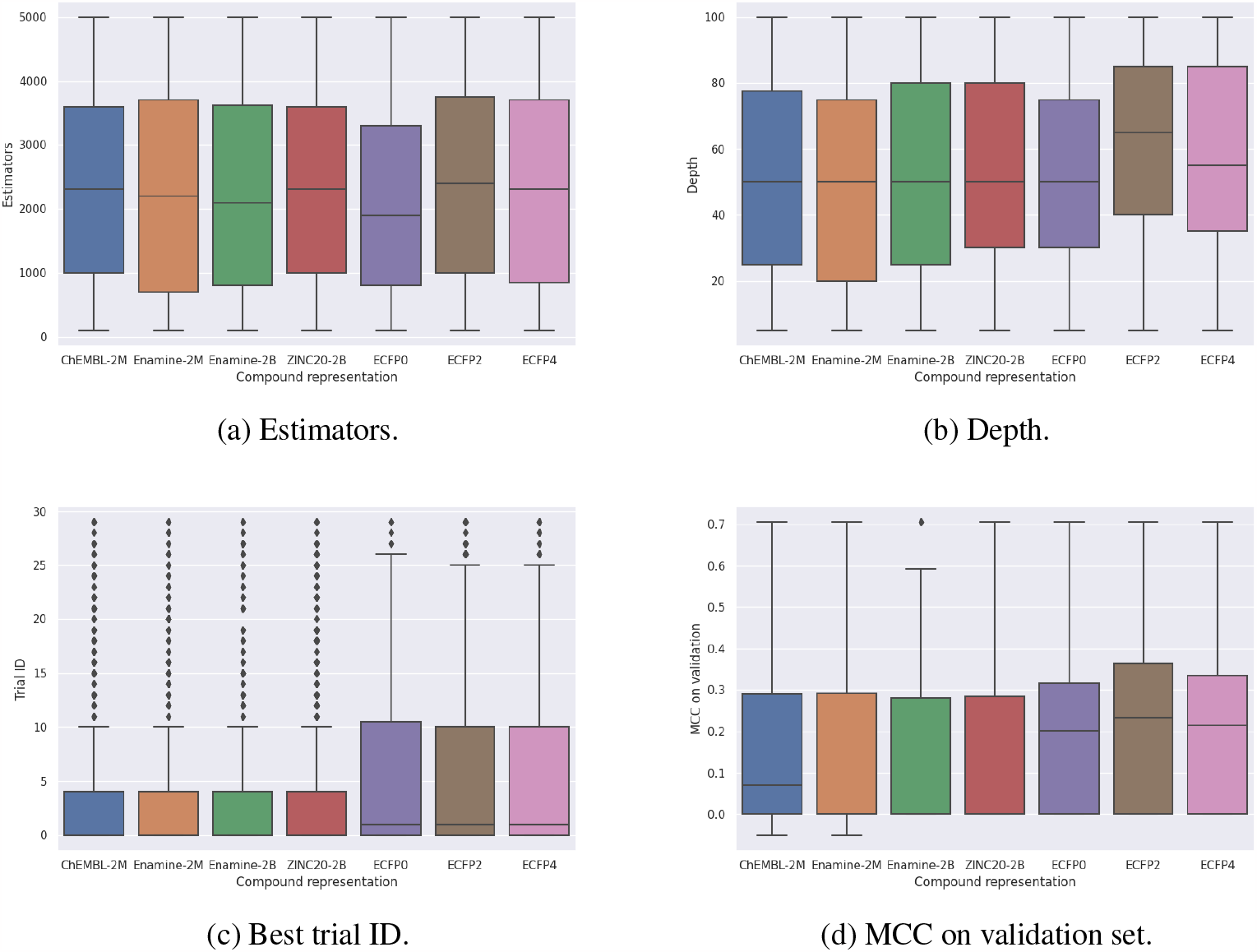
Overview of the best hyperparameters for MoleculeNet - physiology (a-b), the number of trials to find them (c), and the resulting MCC on validation (d).

**Figure 7.**
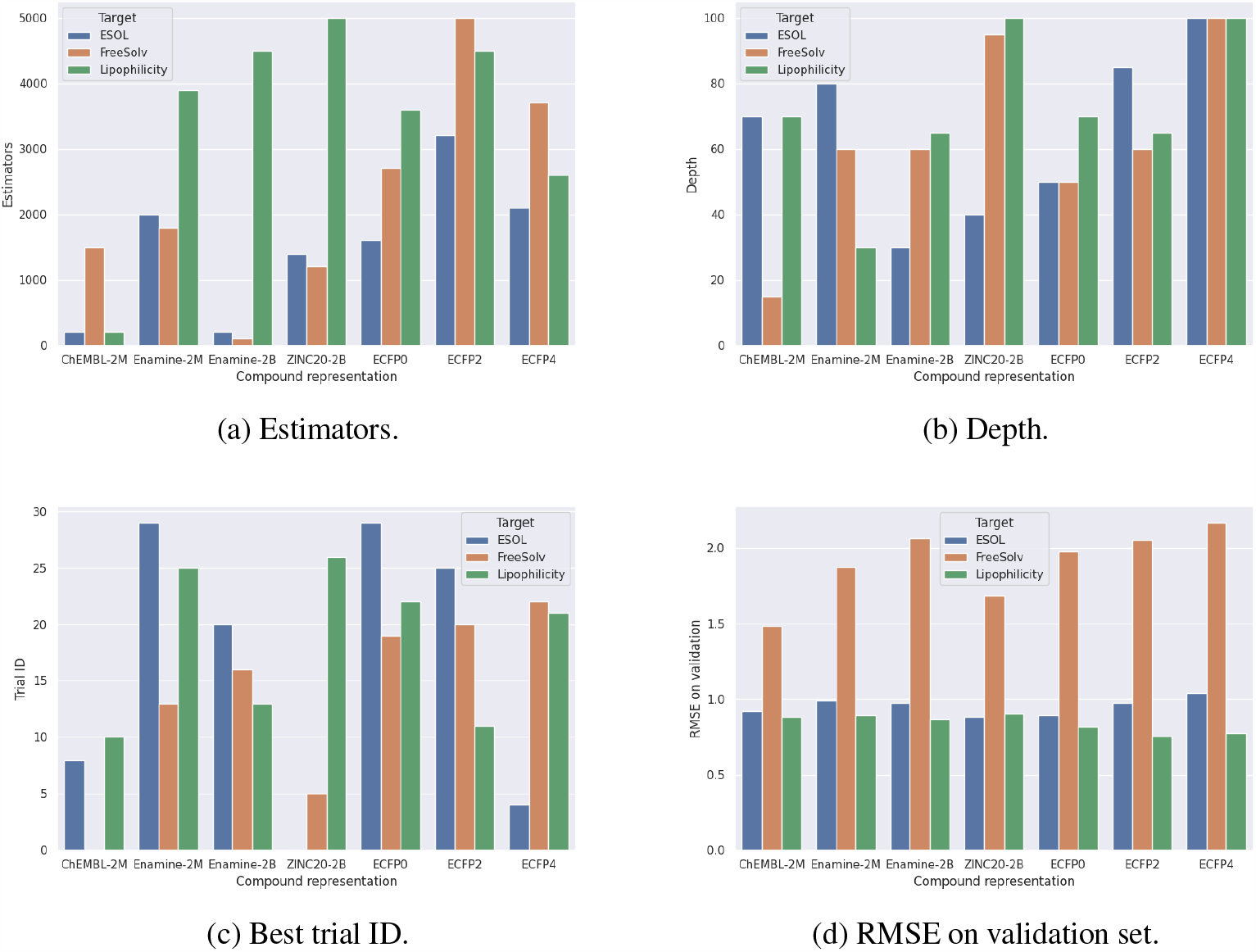
Overview of the best hyperparameters for MoleculeNet - physical chemistry (a-b), the number of trials to find them (c), and the resulting RMSE on validation (d).

